# Regulatory divergences in dosage compensation cause hybrid male inviability in *Caenorhabditis*

**DOI:** 10.1101/2024.02.06.577000

**Authors:** Yongbin Li, Yimeng Gao, Jiaonv Ma, Yifan Gao, Wangyan Zhou, Hantang Zhang, Wenhua Shao, Zhijin Liu, Zhongying Zhao, Xiao Liu

**Author notes:** These authors contributed equally to this work (Yongbin Li, Yimeng Gao). **One-Sentence Summary:** Aberrant initiation of X-chromosome dosage compensation in male embryos underlies asymmetric hybrid male lethality in worms.

## Abstract

The genetic basis of Haldane’s rule, such as hybrid male incompatibility in XX systems, has long remained elusive. Here, we found that crosses of *Caenorhabditis nigoni* males with *C. briggsae* females result in insufficient expression of *Cbr-xol-1*, an X-linked master switch responsible for sex determination, consequently activating aberrant dosage compensation in males, and ultimately leading to embryonic inviability. Three compensatory divergences result in comparable *xol-1* expression levels between the parental species but lethal *Cbr-xol-1* underexpression in hybrid male embryos: 1) a less active *Cbr-xol-1* promoter than its *C. nigoni* ortholog; 2) loss of an X-linked *xol-1* paralog in *C. briggsae*; and 3) pseudogenization of a *C. briggsae* autosomal repressor of *xol-1*. Our results define an evolutionary scenario of sexual incompatibility leading to hybrid male inviability.

## Main Text

One of the few well-accepted generalizations about speciation is Haldane′s rule:“When in the F1 offspring of two different animal races one sex is absent, rare, or sterile, that sex is the heterozygous sex” (*1–4*). While genetic models have been proposed to explain hybrid male incompatibility, the molecular processes that are disrupted in hybrid males remain largely elusive(*1, 5*). A breakdown in dosage compensation has been proposed to cause hybrid male lethality(*1, 6, 7*), although studies in fruitfly and mosquito suggest that dosage compensation cannot explain Haldane’s rule(*8–12*).

In organisms with strongly differentiated sex chromosomes, dosage compensation is an essential regulatory process in which the expression in genes on sex chromosomes is equalized between the two sexes(*13*). Chromosomal sex determination involves an XO system in *Caenorhabditis*, wherein males are the heterogametic sex(*14*). In *C. elegans*, dosage compensation occurs in XX animals to reduce X-chromosome transcription by half (*15*). The master sex-determination switch gene *xol-1* is specifically expressed in XO worms to direct male development and prevent dosage compensation, thereby balancing X expression between XX and XO embryos (*15*). Dosage compensation and sex determination pathways are so conserved across *Caenorhabditis* that *xol-1* mutations can aberrantly initiate dosage compensation in the single X of XO embryos, consequently inducing male lethality in both *C. elegans* and *C. briggsae* species (*13, 16*), despite 15-30 million years (MYR) of evolutionary divergence (*17*),.

By contrast, the sister species *C. briggsae* and *C. nigoni*, diverged approximately 3.7 MYR ago (*18*), and the results of their hybridization are typical of Haldane’s rule(*19*). Interspecific crosses in which *C*. *nigoni* serves as the paternal parent produces only hybrid females(*19*). These unisexual broods are the result of hybrid male inviability, which manifests during embryogenesis(*20*). However, the reciprocal cross produces only slightly female-biased broods after embryo hatching(*20*). This asymmetric hybrid male incompatibility in reciprocal crosses is common in diverse organisms, a phenomenon termed Darwin’s corollary to Haldane’s rule.(*21, 22*).

Here, we found that the expression level of X-chromosome in inviable hybrid male embryos of *C*. *nigoni* (♂) x *C. briggsae* (♀) crosses is half that of X-chromosome expression in wild-type *C. briggsae* males. This dysregulated dosage compensation arises from a six-fold reduction in *Cbr-xol-1* expression in the hybrid males compared to that in the parental species. This insufficient transcription of *Cbr-xol-1* is the result of negative epistatic interactions between X-linked and autosomal loci. More specifically, our analyses showed that the *C. briggsae* X-chromosome lacks a *xol-1* paralog that is functional in *C. nigoni* and the *Cbr-xol-1* promoter is less active than its *C. nigoni* ortholog. In addition, genes in the right arm of *C. nigoni* ChrII dominantly repress *Cbr-xol-1* expression in the hybrid males, one of which is a paralog of the *xol-1*-repressing transcription factor, *sex-1*, which is pseudogenized in *C. briggsae*. In short, our results illustrate a subset of co-evolved compensatory mechanisms responsible for regulating *xol-1* that cause embryonic inviability in hybrid males.

### Dosage compensation occurs as an aberration in inviable hybrid male embryos

To investigate whether hybrid male inviability accumulates progressively during embryogenesis or is restricted to specific stages, we conducted reciprocal crosses between *C. briggsae* and *C. nigoni* (Fig. 1A). Embryos were harvested more than 15 hours after laying to allow the completion of embryogenesis for viable hybrids (Fig. 1B). Consistent with previous report (*20*), very few hybrid male embryos carrying a *C. briggsae* X-chromosome (*Cbr*X) successfully hatched (Fig. 1B). Moreover, the distribution of stages in which arrest occurred in such male embryos was uneven throughout embryogenesis. For instance, 35% of the male embryos arrested within the short developmental window between the 400-cell and bean stages (Fig. 1B).

**Fig. 1.**
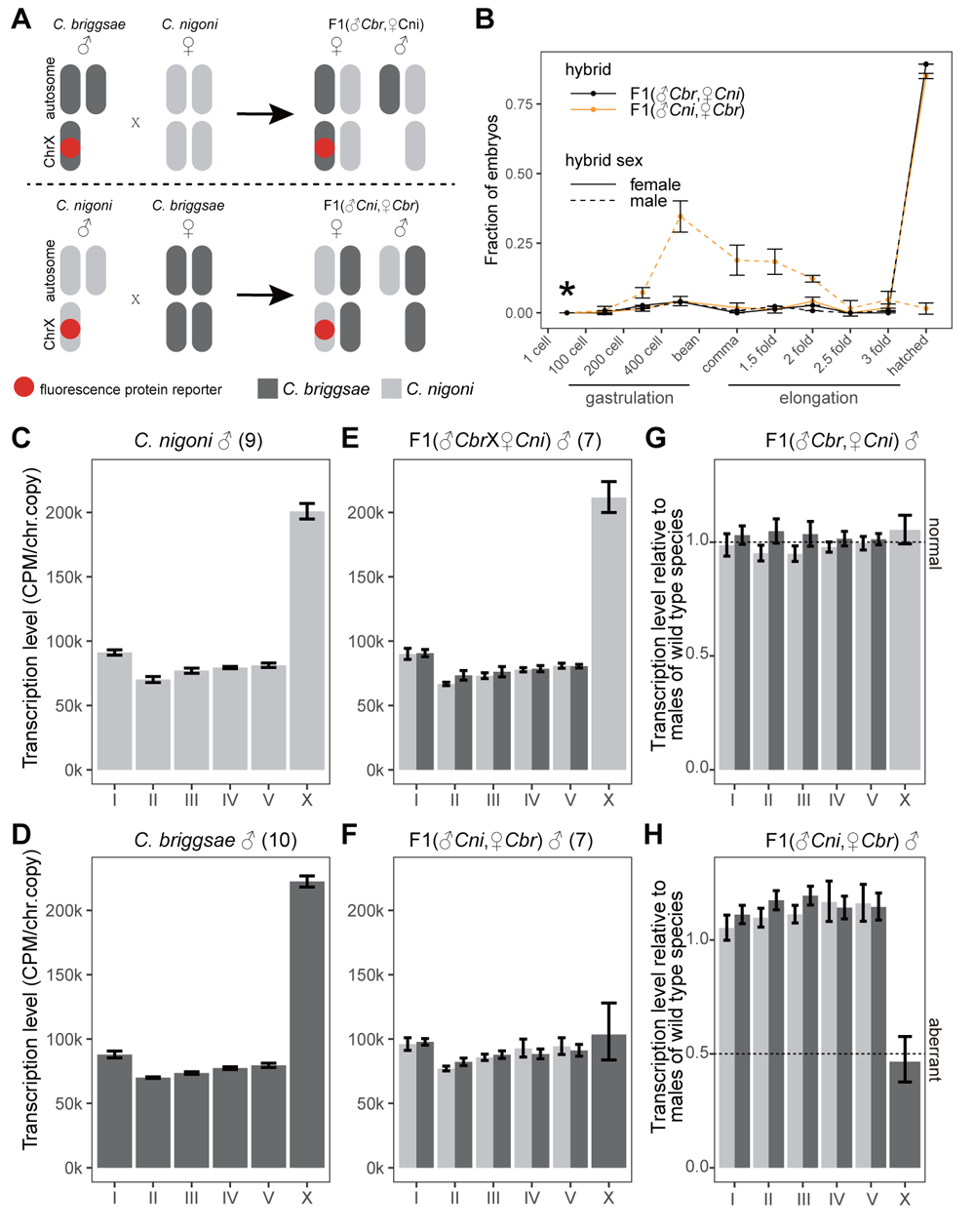
Profiling hybrid embryos from reciprocal crosses between *C. briggsae* and *C. nigoni*. **(A)** Schematic for discerning the sex of hybrid embryos by a visible X-linked marker. **(B)** The proportion of hatched embryos and arrested embryos at each stage across all scored animals for each combination of cross type and sex. Data points represent the mean of three biological replicates. There were very few F1 embryos arrested younger than the 100-cell stage, regardless of cross direction. Embryos were cultured in M9 buffer. (**C** to **F**) Transcription levels of chromosomes in male embryos of *C. nigoni* (C), *C. briggsae* (D), hybrids from *C. nigoni* mothers (E), and hybrids from *C. briggsae* mothers (F). The number of profiled embryos is indicated in parentheses. (**G** and **H**) Ratios of chromosomal transcription levels of hybrid males from *C. nigoni* mothers (G) and hybrid males from *C. briggsae* mothers (H) to males of parental species. Aberrant in (H) denotes an aberrant dosage compensation phenotype. The color codes for species origin of chromosomes are as indicated. error bars, 95% confidence interval.

Since dosage compensation specifically occurs in females to reduce transcription of ChrX by half in order to balance X-linked gene expression between males and females (*15*), we next profiled the transcriptomes of male and female comma stage single embryos, quantifying chromosomal mRNA levels as the sum of Counts per million mapped read (CPM) of all genes in that chromosome. In either parental species, we found no significant difference in the mRNA levels of any chromosome between males and females (fig. S1, A to C and E to G). To account for chromosome dosage, the transcription level of a chromosome was estimated by dividing its mRNA level, as determined above, by the copy number of that chromosome. The female-to-male ratios for autosomal transcription levels were 1, while the ratio for ChrX transcription level was 0.5 in both species (fig. S1, D and H). These results indicated that our single-embryo RNA-seq assay could accurately detect dosage compensation.

Examination of hybrid female embryos showed that the transcription levels of all chromosomes, including ChrX, was indistinguishable between hybrids and either parental species (Fig. S2), thus illustrating the occurrence of dosage compensation in hybrid females. *C. nigoni* males and hybrid males carrying a *C. nigoni* X-chromosome (*Cni*X) did not exhibit significant difference in *Cni*X transcription level (Fig. 1, C, E, and G). So dosage compensation did not occur in *Cni*X-carrying hybrid males, akin to the parental species and consistent with their viability. By contrast, in hybrid males produced by reciprocal cross, *Cbr*X transcription levels were half that observed in *C. briggsae* males (Fig. 1, D, F, and H), indicating an aberrant dosage compensation phenotype in *Cbr*X-carrying hybrid males. Since reduced ChrX expression due to *xol-1* deletion is well-established to result in male lethality in either *C. elegans* or *C. briggsae* (*16, 23, 24*), we therefore hypothesized that the aberrant occurrence (i.e., dysregulation) of dosage compensation was likely the cause of inviability in *Cbr*X-carrying hybrid males.

### Dysregulation of dosage compensation requires CbrX-linked trans-acting factors

The initiation of dosage compensation involves loading the *trans*-acting Dosage Compensation Complex (DCC) onto numerous *cis*-acting sites along ChrX (*15*). To investigate whether dysregulation by *trans*-acting factors or pan-X *cis*-acting sequence elements were responsible for the occurrence of dosage compensation in *Cbr*X-carrying hybrid males, we conducted RNA-seq in male embryos generated by crossing *C. briggsae* males with females of the previously published ZZY10330 or ZZY10337 hybrid introgression lines (HILs). These HILs carry the complete *C. nigoni* genome except for their *Cni*X, which respectively harbor introgressed 5 Mb and 10 Mb fragments of *Cbr*X, and they produce viable hybrid males (Fig. 2, A and B)(*25*). RNA-seq analysis in male progeny of these crosses indicated that transcription levels of each *Cbr*X region (i.e., containing the 10330 or 10337 introgression fragments) were indistinguishable between the progeny and the *C. briggsae* parent (Fig. 2, C and D). These results suggested that dosage compensation did not occur in hybrid males carrying either of these *Cbr*X introgression fragments, and therefore *trans*-acting factors encoded in *C. briggsae* genome outside of the 10330 and 10337 regions was required for dosage compensation-related inviability in hybrid males.

**Fig. 2.**
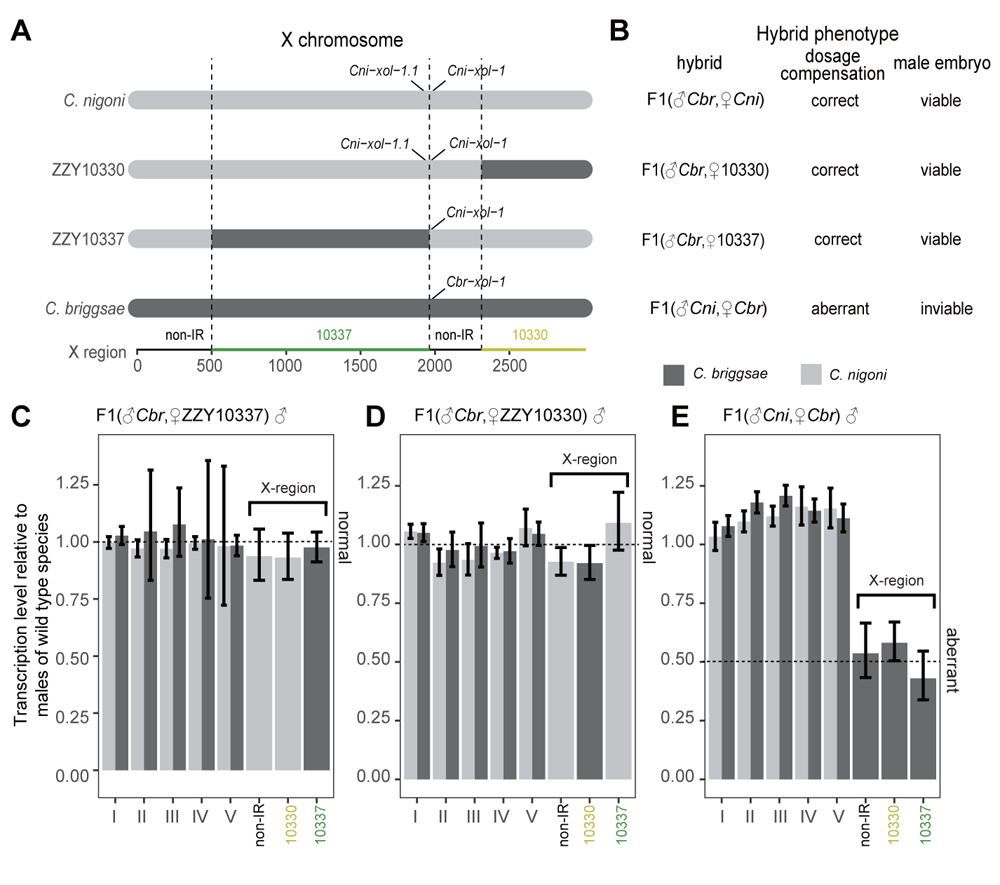
Dosage compensation in hybrid embryos from *C. nigoni* HIL mothers. **(A)** Computational division of ChrX into three X regions. Dark grey horizontal bars in HIL strains ZZY10330 and ZZY10337 represent introgressed *Cbr*X fragments. Dashed vertical lines denote breakpoints between *Cni*X and introgressed *Cbr*X fragments in an HIL strain. The locations of *xol-1* ortholog and paralog are depicted. Orders of orthologous genes along ChrX serve as chromosome coordinates and are indicated below X region. **(B)** Summary of phenotyping data from Fig. 1B and this figure. (**C** and **D**) Ratios of chromosomal transcription levels of hybrid male embryos from ZZY10337 mothers (C), and hybrid male embryos from ZZY10330 mothers (D) to male embryos of two native species. Relative chromosomal transcription levels are calculated using seRNA-seq data shown in fig. S3. (**E**) Ratios of chromosomal transcription levels of hybrid male embryos from *C. briggsae* mothers to male embryos of parental species. Same seRNA-seq data were used as in Fig. 1H except that all three *Cbr*X regions were calculated together in Fig. 1H. The color codes for species origin of chromosomes are as indicated. error bar, 95% confidence interval.

Re-examination of the transcription levels of the 10330 and 10337 regions in male hybrid embryos produced in the original *C. nigoni* (♂) X *C. briggsae* (♀) crosses confirmed that these regions were each subjected to dosage compensation (Fig. 2E). Since the viable males (without dosage compensation) produced by *C. briggsae* (♂) X 10330 or 10337 (♀) crosses carried *Cni*X non-introgressed regions (non-IRs), while embryonic inviable hybrid males carried the *Cbr*X non-IRs (Fig. 2B), it was reasonable to conclude that the relevant trans-acting factors were likely located within these *Cbr*X non-IRs.

### C. briggsae lost an X-linked xol-1 paralog

We searched these *Cbr*X non-IRs for orthologs of the ten DCC subunits and its six upstream regulators identified in *C. elegans* (*15*) (Fig. 3A). We found one such candidate ortholog, *xol-1*, located in the *Cbr*X right non-IR (Fig. 2A). In *C. briggsae, Cbr-xol-1* reportedly functions as a master switch to negatively regulate dosage compensation in males and is therefore necessary for male viability (*16*). The *C. nigoni* genome codes two copies of *xol-1, Cnig_chr_X.g24950* and *Cnig_chr_X.g24897* (*18, 26*). As an *xol-1* ortholog*, Cnig_chr_X.g24950* was annotated as *Cni-xol-1* (fig. S4A). Positioned 0.3 Mb upstream of *Cni-xol-1*, *Cnig_chr_X.g24897* was identified as an *xol-1* paralog, resulting in its annotation as *Cni-xol-1.1* (Fig. 2A).

**Fig. 3.**
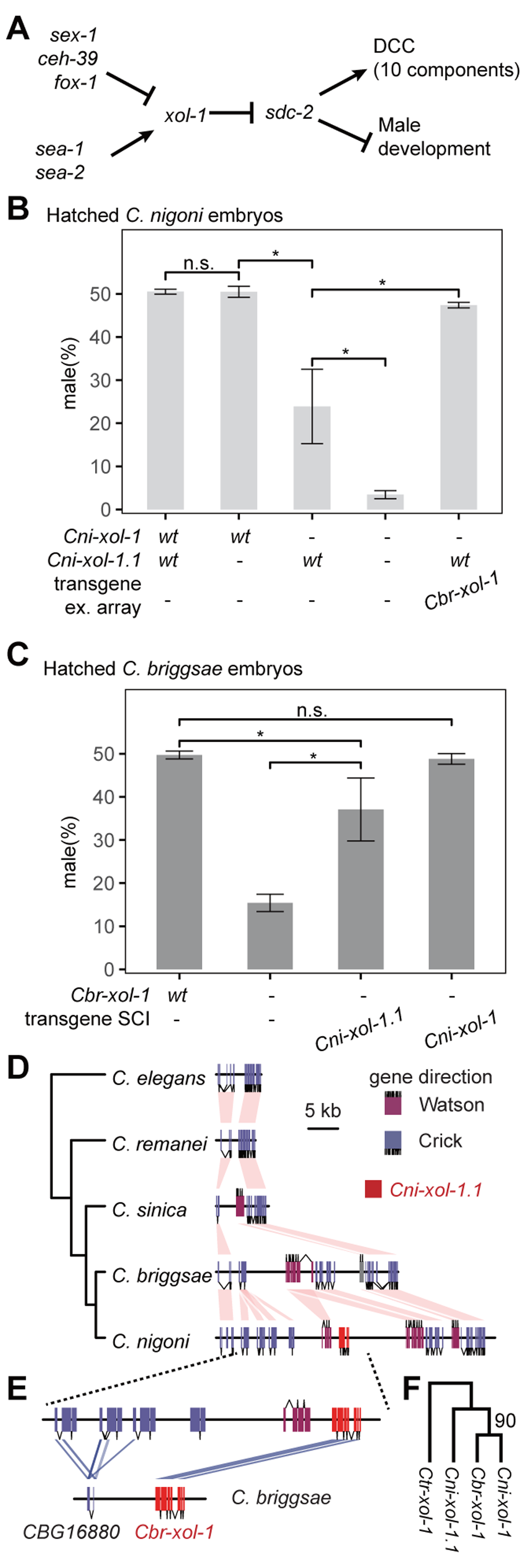
Interspecific divergence of *xol-1* in copy number. **(A)** Overview of regulatory hierarchy that controls sex determination and dosage compensation in *C. elegans*. Dosage Compensation Complex component genes: *capg-1, dpy-21, dpy-26, dpy-27, dpy-28, dpy-30, mix-1, sdc-1,* and *sdc-3* as well as *sdc-2*. (**B** and **C**) The fraction of males among hatched embryos of *C. nigoni* (B) and *C. briggsae* (C). Statistical significance is based on Student’s *t* test; n.s., not significant;,error bars denote standard deviation.; and * indicates *p* < 0.05, ** indicates *p* ≤ 0.01, and *** indicates *p* ≤ 0.001. (**D**) A *Cni*X segment where *Cni-xol-1.1* resides and its orthologous blocks in other five *Caenorhabditis* species. Pink pastel shapes connect orthologous genes. (**E**) Synteny between a *Cni*X block where *Cni-xol-1.1* resides and a *Cbr*X block where *Cbr*-*xol-1* resides. The *Cbr*X block is reverse complementary to the reference genome. Blue pastel shapes connect homologous sequences, e.g., *CBG16880* and its three most homologous *C. nigoni* genes based on tBLASTn. (**F**) Phylogenic tree of *xol-1* orthologs and *Cni-xol-1.1*. Out group is C*. tribulationis xol-1* ortholog (*Ctr-xol-1*). The percentage of replicate trees in which the associated taxa clustered together in the bootstrap test is shown next to the branch if it is > 80.

Disruption of *Cni-xol-1.1* had no significant effect on hatching rates in male *C. nigoni* embryos, while mutation of *Cni-xol-1* resulted in a significant decrease in male hatching rate (Fig. 3B). Moreover, double mutation of *Cni-xol-1* and *Cni-xol-1.1* further decreased male embryo viability (Fig. 3B). These results indicated that *Cni*-*xol-1* and *Cni*-*xol-1.1* function redundantly in embryogenesis, with *Cni-xol-1* having stronger activity. To assess whether *xol-1* genes are functionally conserved between the sister species, we introduced a multi-copy *Cbr-xol-1* extra-chromosomal array transgene (ex[*Cbr-xol-1*]) into *C. nigoni* worms knocked out for *Cni-xol-1* and found that the ortholog could likewise rescue the impaired viability of male embryos (Fig. 3B). Subsequently, we used Cas9-mediated Single Copy Insertion (Cas9-SCI) transgenesis to integrate each *C. nigoni xol-1* gene into the *C. briggsae* genome and referred to these two transgenes as SCI[*Cni-xol-1*] and SCI[*Cni-xol-1.1*], respectively. While SCI[*Cni-xol-1*] could fully rescue inviability in *Cbr-xol-1* null male embryos, SCI[*Cni-xol-1.1*] could only partially rescue this phenotype (Fig. 3C). These reciprocal rescue assays demonstrated the conserved function of these three *xol-1* genes in determining male embryo viability. Nevertheless, the paralog, *Cni*-*xol-1.1,* had weaker activity than both *xol-1* orthologs.

Synteny analysis showed that *Cni-xol-1.1* resides in a ChrX region unique to both *C. briggsae* and *C. nigoni,* and all genes except *Cni-xol-1.1* in this unique ChrX region have syntenic orthologs between the two species (Fig. 3D). Moreover, this region in *C. nigoni* carries all of the top-three homologous copies of the *Cbr-xol-1* downstream gene, *CBG16880*, which are arranged as tandem repeats downstream of *Cni-xol-1.1* (Fig. 3E). This synteny supports the possibility that *Cni-xol-1.1* originated from a segmental duplication involving the ancestral *xol-1* and *CBG16880* prior to the divergence of these sister species. Moreover, phylogenic analysis of *xol-1* genes showed a significant branching of *Cni-xol-1.1* from the *C. briggsae* and *C. nigoni xol-1* ortholog pair (Fig. 3F)*. Together, these* synteny and phylogenetic analyses suggest that the segmental duplication which gave rise to *xol-1.1* likely occurred in the common ancestor of *C. briggsae* and *C. nigoni*, after which, *xol-1.1* was lost in *C. briggsae* following divergence from *C. nigoni*.

### Low expression of Cbr-xol-1 causes hybrid male embryo inviability

To assess whether replacing *Cbr-xol-1* with *C. nigoni xol-1* genes could rescue hybrid male inviability, we crossed *C. nigoni* males to *Cbr-xol-1* knockout *C. briggsae* females carrying the SCI[*Cni-xol-1*], the SCI[*Cni-xol-1.1*], or both. Hybrid male embryos only carrying the SCI[*Cni-xol-1.1*] still had a small hatching rate, whereas those only carrying the SCI[*Cni-xol-1*] showed significant increase in viability (Fig. 4A). The SCI[*Cni-xol-1*] and the SCI[*Cni-xol-1.1*] together could further improve the hatching rate of hybrid male embryos to a level indistinguishable from the *Cni*X-carrying positive control hybrid males (Fig. 4A). To examine if the high rescue capability of the double *xol-1* combination depends on species origin of *xol-1* ortholog, we crossed *C. nigoni* males to *C. briggsae* females carrying endogenous *Cbr-xol-1* and the SCI[*Cni-xol-1.1*]. We found that hybrid male embryos from this cross had a hatching rate even lower than wild-type hybrid males (Fig. 4A). The dramatic difference in rescue capability between these two *xol-1* orthologs suggested lower gene activity of *Cbr-xol-1* than *Cni-xol-1*.

**Fig. 4.**
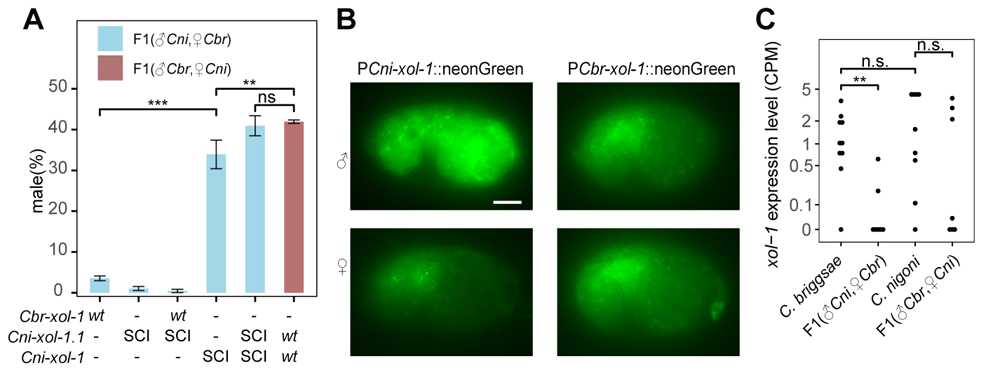
Association of *xol-1* expression level with hybrid male embryo viability. (**A**) The fraction of males among hatched hybrid embryos. SCI, the Single Copy Insertion alleles depicted in Fig. 3C. (**B**) Expression patterns of promoter reporters of *xol-1* orthologs in *C. briggsae* embryos. Embryos were photographed at bean stage and with same exposure time. In photograph, the anterior end of each embryo was arranged to right side and ventral to bottom. Putative signal in female embryos and SCI[PCbr-xol-1::neonGreen] males is gut autofluorescence. Scale bars, 10 μm. **(C)** The *xol-1* expression level quantified by seRNA-seq. The expression levels of *Cni-xol-1* and *Cni-xol-1.1* are combined because some of their reads are indistinguishable. A dot represents a male embryo. This dot plot is based on same smRNA-seq data as those used in Fig. 1 and fig. S1.

To compare the promoter activity of the *xol-1* orthologs, we constructed *Cbr-xol-1* and *Cni-xol-1* promoter fusion reporters. Using Cas9-SCI transgenesis, we integrated single copies of each reporter at the same site as the SCI[*Cni-xol-1*] in the *C. briggsae* genome. The *Cni-xol-1* promoter reporter displayed significant male-specific expression in *C. briggsae* embryos, while signal from the *Cbr-xol-1* promoter was undetectable (Fig. 4B), indicating that *Cni-xol-1* was expressed at higher levels than its ortholog in *C. briggsae* under the same trans environment. Subsequently, we compared *xol-1* expression levels in male embryos between parental species and hybrids by further analysis of the single-embryo RNA-seq data. We observed that *xol-1* expression levels were similar among viable *Cni*X-carrying hybrids, *C. nigoni*, and *C. briggsae*, but significantly lower in inviable *Cbr*X-carrying hybrids (Fig. 4C). These results supported that interspecific divergences in *xol-1* gene number and expression level of *xol-1* ortholog contribute to hybrid male embryo inviability.

Employing single-molecule inexpensive fluorescence *in situ* hybridization (smiFISH)(*27*), we measured *xol-1* expression levels in embryos from the 500-cell through bean stages, which were enriched for embryonic arrest in *Cbr*X-carrying hybrid males (See Fig. 1B). In these stages, both parental species and hybrids displayed *xol-1* expression in males ( Fig. 5, A, B, D, and F) higher than in females (fig. S5). The abundance of cytoplasmic *xol-1* mRNA puncta did not significantly differ between male embryos of *C. nigoni* and those of *Cni*X-carrying hybrids (Fig. 5, A to C), which were viable. In contrast, the abundance of cytoplasmic *Cbr-xol-1* mRNA signals in inviable *Cbr*X-carrying hybrid males was only one sixth that in *C. briggsae* males and the residual *Cbr-xol-1* signal was localized to the posterior regions of embryos (Fig. 5, D, F and H). To validate the effect of *xol-1* transcription level on viability of hybrid male embryos, we constructed three respective expression vectors for *Cbr-xol-1*, *Cni-xol-1*, or *Cni-xol-1.1* driven by the constitutive promoter, *Cel-eft-3*. We found that injecting each vector into *C. briggsae,* separately, resulted in significant rescue of *Cbr*X-carrying hybrid male embryos (Fig. 5I). These collective results suggested that insufficient *xol-1* transcription was a cause of hybrid male embryo inviability.

**Fig. 5.**
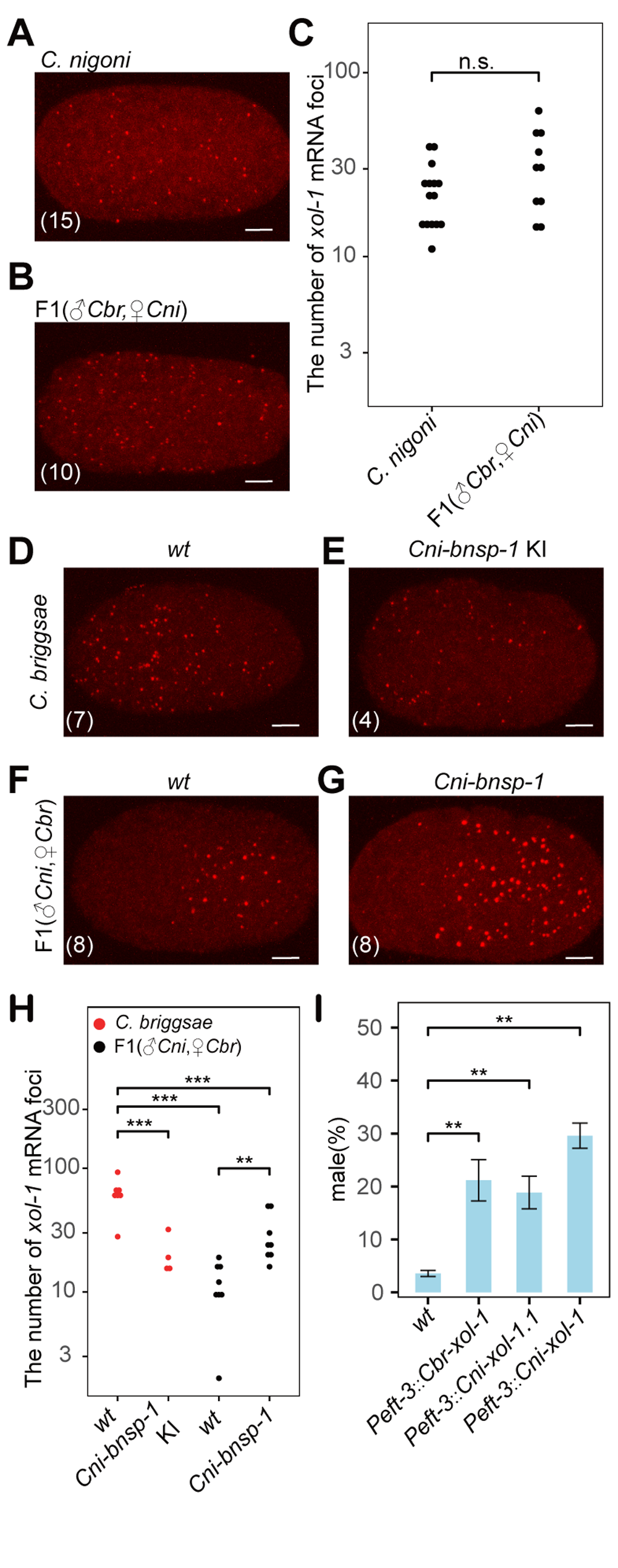
The mRNA molecules of *xol-1* genes detected in male embryos by smiFISH. **(A** and **B)** Cytoplasmic *Cni-xol-1* and *Cni-xol-1.1* mRNA molecules in male embryos of *wild-type C. nigoni* (A) and hybrids sired by *wild-type C. briggsae* (B). Probes used in these embryos are designed for both *C. nigoni xol-1* genes so that a punctum can be either *Cni-xol-1* mRNA or *Cni-xol-1.1* mRNA. Every embryo is oriented so that its anterior region is to the left. The number of scored embryos is indicated in parentheses. Embryos were collected from the 500-cell to the bean stages. **(C)** Statistics of *Cni-xol-1*/*Cni-xol-1.1* mRNA abundance in (A) and (B). Each dot in the plot represents an embryo. The logarithm of the number of *xol-1* puncta was used for Student’s *t* test. **(D** to **G)** Cytoplasmic *Cbr-xol-1* mRNA molecules in male embryos of *wild-type* (D) and *Cni-bnsp-1* Knock-in *C. briggsae* (E), as well as hybrids sired by *wild-type* (F) and *Cni-bnsp-1* null *C. nigoni* (G). Probes used in these embryos are designed for *Cbr-xol-1.* Embryos were processed and photographed as in (A) and (B). **(H)** Statistics of *Cbr-xol-1* mRNA abundance data in (D) to (G). (**I**) The fraction of males among hatched hybrid embryos carrying *Cbr*X and *xol-1* transgenes driven by housekeeping promoter *Cel-eft-3*.

### Evolution of an autosomal xol-1 repressor contributes to hybrid male inviability

As F1 male progeny of *C. briggsae* (♂) X *C. nigoni* ZZY10337 (♀) crosses are fertile(*28*), we can use these F1 males to screen for specific ChrX-autosome interactions that might contribute to hybrid male inviability (Fig. 6A). To this end, we crossed fertile F1 males with *C. briggsae* hermaphrodites, then collected the resulting viable B2 males for genotyping by single-worm genome sequencing (Fig. 6B). We expected that autosomal loci in *C. nigoni* that could suppress hybrid male viability should be underrepresented among these viable B2 males. Analysis of sequencing data identified right part of *Cni*II (position 12.694Mb to right end) as the most underrepresented locus, and underrepresentation of this *Cni*II locus was male-specific, not occurring in viable B2 females (Fig. 6C). Furthermore, this interval overlapped with an interval of *Cni*II previously found to contribute to hybrid male inviability(*28*) (Fig. 6C). By focusing on the overlapping segments between our data and the previous report, we could further narrow the candidate region of *Cni*II to approximately 1.8 Mb.

**Fig. 6.**
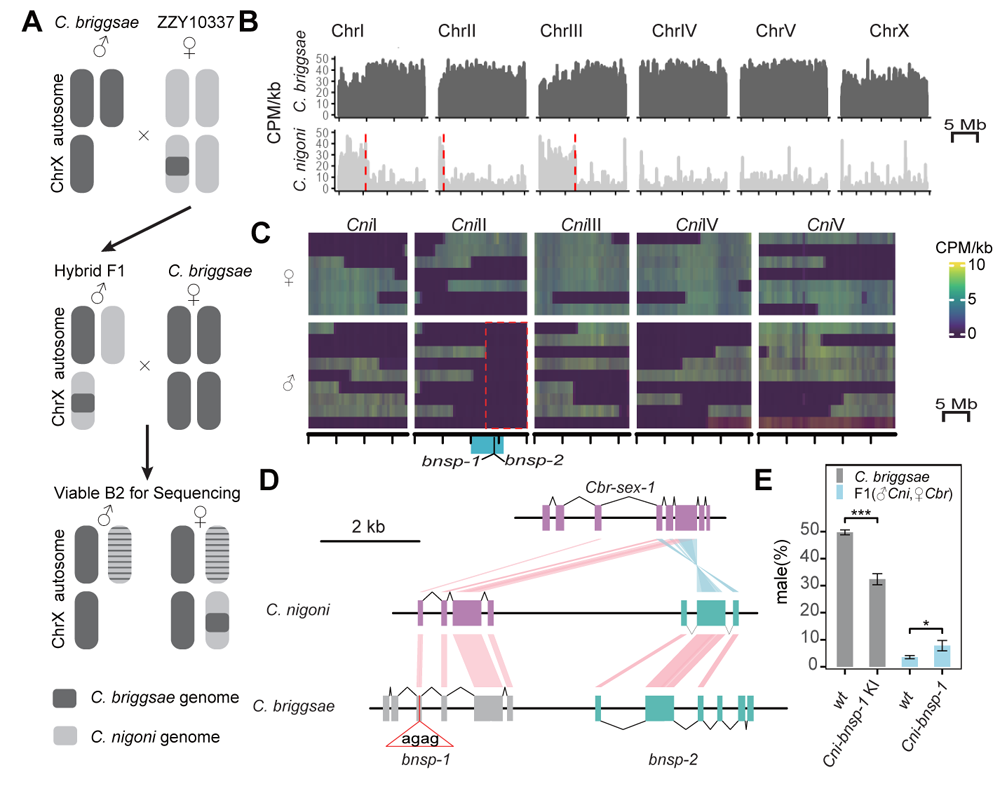
Evolution of *bnsp-1* and its role in hybrid male inviability. **(A**) Backcrossing strategy to screen for *C. nigoni* autosomal loci dominantly suppressing hybrid male viability. **(B)** DNA-seq data of a viable B2 male adult. Every recombinant autosome from an F1 male is expected to carry only one recombination site, which is delineated by a vertical dashed line. **(C)** The heatmap of *C. nigoni* autosomal reads in viable B2 males and females. The deduced smallest interval responsible for hybrid male inviability is highlighted by a red dashed box. The blue bar below the *Cni*II indicates previously mapped autosomal interval repressing hybrid male viability (see main text). The locations of two se*x-1* paralogs, *bnsp-1* and *bnsp-2*, are depicted. **(D)** Interspecific divergences in *bnsp-1* and *bnsp-2* genes. *Cbr*-*bnsp-1* is in grey because it is a pseudogene. The 4-bp insertion in *Cbr*-*bnsp-1* is depicted. (**E**) The fraction of males among hatched embryos.

Notably, no gene in this region is orthologous to any of the 16 genes involved in sex determination and dosage compensation in *C. elegans* (Fig. 3A). However, this chromosomal segment contained two paralogs of the X-linked sex determinant, *sex-1* (Fig. 6C). In *C. elegans*, *sex-1* encodes a nuclear hormone receptor that binds to and represses the *xol-1* promoter in XX embryos, while mutation of *sex-1* can enhance *xol-1* transcription and consequently XX lethality (*15*). These two *sex-1* paralogs were only present in the *C. briggsae*/*C. nigoni* branch of *Caenorhabditis* (fig. S4B), leading to their annotation as *C. <u>b</u>riggsae*/*C. <u>n</u>igoni <u>s</u>ex-1 <u>p</u>aralog-1/2* (*bnsp-1* and *bnsp-2*, respectively; Fig. 6D). Their most striking interspecific divergence is pseudogenization of *Cbr-bnsp-1* resulting from a 4-bp insertion in exon 2, whereas its *C. nigoni* ortholog has an intact open reading frame (Fig. 6D).

To test whether *Cbr-bnsp-1* can be regulated by *Cni*-*bnsp-1*, we generated a *C. briggsae* strain with pseudogene *Cbr-bnsp-1* replaced by *Cni-nsp-1* via CAS9-mediated Knock-in. Subsequent smiFISH analyses showed that *Cbr*-*xol-1* expression in male embryos significantly decreased in the KI[*Cni-nsp-1*] strain (Fig. 5, D, E and H), indicating that *Cni-bnsp-1* represses *Cbr*-*xol-1* expression. Consistent with observed *Cbr*-*xol-1* downregulation, this KI[*Cni-nsp-1*] strain showed significant reduction in the hatching rate of male embryos (Fig. 6E). Next, we generated an *Cni*- *nsp-1* null *C. nigoni* strain using CAS9-mediated Knock-out. Hybrid male embryos sired by the *Cni-nsp-1* null strain had significantly higher *Cbr-xol-1* expression than those sired by the wild-type *C. nigoni* strain, but still significantly lower than *C. briggsae* male embryos (Fig 5, D and F to H). Consistent with the partial *Cbr*-*xol-1* upregulation, disruption of *Cni-nsp-1* significantly rescued *Cbr*X-carrying hybrid male embryos, but only to a small degree (Fig. 6E). These results thus demonstrated that *Cni-nsp-1* functioned as a repressor of *Cbr-xol-1*, contributing to hybrid male embryo inviability with other trans-acting factors in the right arm of *Cni*II.

## DISCUSSION

This study of the *C. briggsae* and *C. nigoni* sister species showed that hybrid male inviability is caused by insufficient transcription of *xol-1*, an X-linked master gene required for sex determination and the dosage compensation pathway (*15, 16*). Although *C. briggsae* maintains *xol-1* expression level comparable to that of *C. nigoni* through compensatory *cis-trans* regulatory divergence, divergence in *cis*-acting elements results in a relatively weaker *Cbr-xol-1* promoter than the *Cni-xol-1* promoter. Moreover, the *C. briggsae* genome lacks a *xol-1* paralog present in *C. nigoni*. Alternatively, the *C. nigoni* genome encodes a functional *Cbr*-*xol-1*-repressing transcription factor, *bnsp-1*, whereas *C. briggsae* carries only a pseudogenized ortholog. By contrast, hybrid male progeny of crosses between *C. briggsae* males and *C. nigoni* females carry both the *xol-1* paralog and the strong promoter of *xol-1* ortholog, and as a result, express *xol-1* at levels comparable to the parental species. In hybrid males generated by reciprocal cross, *Cbr-xol-1* with a weak promoter is both hemizygous and repressed by functional *Cni*-*bnsp-1*. This combination of *cis-* and *trans-* factors leads to insufficient *xol-1* expression, dysregulated dosage compensation, and ultimately hybrid male embryo inviability.

In *Caenorhabditis*, the sex determination cascade is initiated by relaying the ratio between the number of X chromosomes and sets of autosomes to productive *xol-1* mRNA level (*15, 16, 29*). The activity of *xol-1* then controls the rest of the sex determination pathway(*30*). Findings in this study of *xol-1* and its regulator *nsp-1* illustrate three evolutionary features of the sex determination pathway in different taxa. First, the master regulators of the sex determination pathway tend to change dramatically, while genes at the bottom of the hierarchy remain the relatively conserved between species (*31, 32*). Second, the sex determination pathway is modified through variation, duplication, or loss of sex-determining genes (*33, 34*). Third, DNA and protein sequence of sex determination genes appear to evolve at a rapid pace (*34*). These findings thus highlight the dynamic nature of sex determination and dosage compensation pathway and its susceptibility to disruption in interspecies hybrids.

The ubiquity of Haldane’s rule implies that postzygotic incompatibility evolves through similar mechanisms across a wide range of taxa (*35*). Crosses between the sibling species, *Drosophila melanogaster* and *D. simulans,* result in unisexual lethality. Hybrid males typically die at the larval/pupal transition when *D. melanogaster* is the maternal parent, while females are embryonic lethal from the reciprocal cross (*36*). However, the role of dosage compensation in their hybrid male lethality remains controversial (*8*). In *Drosophila*, X-linked *sxl* is the master sex determination and dosage compensation gene (*37*) and is female-specific (*38*). Ectopic expression of *sxl* in males leads to male lethality(*39*). Inviable hybrid male embryos carrying the *D. melanogaster* X-chromosome aberrantly express *sxl* (*10*). Moreover, mutation of s*xl* increases larval viability in hybrid males(*10*). But not supporting involvement of dosage compensation in hybrid male inviability, mutation of four essential genes in the *D. melanogaster* dosage compensation pathway does not negatively impact hybrid male viability (*8*). Instead, all three genes critical for hybrid male inviability between these two sister *Drosophila* species encode chromatin proteins: *Hmr*, *Lhr* and *gfzf* (*40–42*) and their involvement in dosage compensation has not been reported. Nevertheless, a loss-of-function *Lhr* mutation rescues hybrid male viability and represses aberrant *sxl* expression in byrbid male embryos (*10*). Further investigation may yet uncover whether dosage compensation is also disrupted in hybrid males carrying the D. *melanogaster* X-chromosome.

Through a pair of experimentally tractable sister species that are capable of producing fertile hybrid offspring and allow large-scale crossing experiments, our study depicts the genetic contributions to hybrid incompatibility. As efforts to discover new nematode species accelerate (*43*), an increasing number of species pairs have emerged with partial intrinsic post-zygotic isolation, such as *C. remanei* and *C. latens*, which exhibit asymmetric hybrid male sterility and inviability (*44, 45*). The genetic resources and experimental tools available for nematodes will drive advances in our molecular genetic understanding of the mechanistic basis responsible for reproductive isolation.

In general, this should include a brief (1-2 paragraph) introduction, followed by a statement of the specific scope of the study, followed by results and then interpretations. Please avoid statements of future work, claims of priority, and repetition of conclusions at the end.

## Acknowledgments

The authors thank Xianyong Sheng for from the Equipment Platform, Capital Normal University for confocal imaging. We are grateful to Dr. Lilin Du and Mengqiu Dong from National Institute of Biological Sciences, Beijing, Dr. Qianghua Xia from Tianjing Medical University, Dr. Zhuo Du from Institute of Developmental Biology and Genetics, CAS, Dr. Jian Lu from Peking University, and Dr. Wen Wang from Northwest Polytechnical University for helpful discussions and comments on biological results. The *Caenorhabditis* Genetics Center provided transgenic worms.

## Funding

National Natural Science Foundation of China grant 31861163005 (XL)

National Natural Science Foundation of China grant 31871472 (XL)

National Natural Science Foundation of China grant 32070639 (XL)

R&D Program of Beijing Municipal Education Commission Grant KM202310028008 (YL)

## Author contributions

Conceptualization: XL, YL, YMG

Methodology: YL, YMG, JM, YFG, WZ, HZ, WS, ZL, ZZ

Investigation: XL, YL, YMG

Visualization: YMG, YL

Funding acquisition: XL, YL

Project administration: XL, YL

Supervision: XL

Writing – original draft: XL, YL, YMG

Writing – review & editing: XL, YL

## Competing interests

Authors declare that they have no competing interests.

## Data and materials availability

All data are available in the main text or the supplementary materials. All sequencing data from this study have been submitted to the NCBI Sequence Read Archive (SRA; www.ncbi.nlm.nih.gov/sra) with BioProject accession number PRJNA1068349 and PRJNA1071360.

## Supplementary Materials

Materials and Methods

Figs. S1 to S5

Tables S1 to S5 References (*1–26*)

Data S1 to S3

## Notes

### Competing Interest Statement

The authors have declared no competing interest.

