## Supplementary Materials for "Regulatory divergences in dosage compensation cause hybrid male inviability in *Caenorhabditis*"

**The PDF file includes:**

Materials and Methods

Figs. S1 to S5

Tables S1 to S5

References

**Other Supplementary Materials for this manuscript include the following:**

Data S1 to S3

Materials and Methods

Nematode strains and maintenance

All worms were cultured at 25 ℃ on OP50-seeded nematode growth medium (NGM) plates with 2.5 % concentrated agar, unless otherwise indicated. This enhanced composition of NGM is specifically designed to prevent the penetration of nematodes into the medium, addressing the common issue of worm burrowing of *C. briggsae* and *C. nigoni*. The inbred *wild-type* *C. nigoni* strain JU1421 and *C. briggsae* strain AF16 were obtained from the Caenorhabditis Genetic Center (CGC). Gonochoric *C. briggsae* strain *Cbr-she-1*(*v49*)(*1*) was gifts from Ronald Ellis. All strains used in this study are listed in Table S1.

To select or enrich worms carrying P*rps-27*::NeoR::*unc-54*-UTR transgene , 500 μl of 12.5 mg/ml G418 solution was added to each seeded 60mm NGM plate for long-term culturing. Before each genetic assay, the enriched worms were recovered at least one generation on non-G418 seeded plates.

Molecular Biology

The Cas9 CDS was codon-optimized for *E. coli*, synthesize and cloned to the pET28a expression vector by RuiBiotech (Beijing, China), creating the pET28a-Cas9 vector. This construct is characterized by the inclusion of a 6xHis tag, a thrombin cleavage site, an SV40 Nuclear Localization Signal (NLS), the Cas9 protein, and another 6xHis tag. For the expression of Cas9, the pET28a-Cas9 vector was transformed to BL21(DE3) Chemically Competent Cell purchased from TransGen Biotech (Beijing, China)(#CD601) or T7 Express Competent E. coli (NEB, #C2566H). Recombinant Cas9 protein was prepared as described(*2*).

For knock-in or Single Copy Insertion, homology recombination (HR) templates were constructed by cloning homology arms and insert sequence into pEAZY-Blunt Zero with Seamless Cloning Kit (Beyotime Biotechnology, Shanghai, #D7010M) and transformed to Trans5α Chemically Competent Cell (TransGen Biotech, #CD201) or transformed to Trans5α directly and assembled with DH5a-mediated assembly(*3*). Target sites in the homology arms of HR templates were modified with synonymous mutations to avoid Cas9 digestion.

Microinjection and transgenesis

Transgenic lines were generated by injecting DNA, RNA, and/or CAS9 protein into the gonads of the indicated worm strains as described(*4, 5*). For androdioecious strains, injected were young adult hermaphrodites. For dioecious strains, injections were performed on pregnant young adult females and then mated with males. For extra chromosomal array or transient expression, the concentration of DNA constructs was 50 ng/µl and the co-injected visible marker was either pZZ184 (p*Cbr-myo-2*::Cherry) (*6*) or pZZ31 (p*Cbr-myo-2*::GFP)(*7*). Sequence of injected vectors is listed in Zenodo(*8*).

CRISPR/Cas9-mediated genome editing

The standalone verion of CCTOP(*9, 10*) was used to select the target sequence. We purchased sgRNAs from GenScript Biotech Corporation (Jiangsu, China). Sequences of sgRNAs and genotyping PCR primers are listed in Table S2 and Table S3 respectively. A *dpy-10* Co-CRISPR strategy(*11*) was employed to enrich edited progenies. Dumpy F1 progenies were selected following injections for PCR genotyping to identify successful genome editing events. Subsequent crossings of these edited worms with wild-type counterparts were carried out to eliminate *dpy-10* mutation.

Knock-out

We employed a dual-sgRNA approach to delete targeted sequence. In brief, 5 ug/ul final concentration of the recombinant Cas9 protein was incubated with two distinct sgRNAs, each at a concentration of 24 uM, and *dpy-10* co-CRISPR gRNA at a concentration of 12 uM, at 37 ℃ for 10 minutes proceeding to injection. Sequence of KO alleles is listed in Zenodo(*8*).

Knock-in and Single Copy Insertion (SCI)

Donor vectors were extracted with FastPure Plasmid Mini Kit purchased from Vazyme Biotech (Nanjing, China) (Item # DC201), as well as purified and concentrated with VAHTS DNA Clean Beads purchased from Vazyme Biotech (Nanjing, China) (Item #N411). To facilitate homology-directed repair, donor vectors were administered at a minimum final concentration of 200 ng/µl per injection. Sequence of KI and SCI alleles is listed in Zenodo(*8*).

Cross setups to distinguish selfing from hybrid offspring and to distinguish male from female embryos

In a cross with hermaphrodites as mothers, father worms carried a membrane localized fluorescence reporter inserted into ChrII, thuB120[P*Cbr-his-72*::myri::cherry] *Cbr*II or thuN26[P*Cbr-his-72*::myri::cherry] *Cni*II. Progenies expressing these reporters were considered as hybrids. In a cross with females as mothers, such as *C. nigoni* XX and *Cbr-she-1*(*v49*) *C. briggsae* XX(*1*), females were picked no later than the L4 stage to ensure virginity.

In crosses whose embryonic offspring needed to have their sexes discerned, father worms carried an X-linked fluorescence reporter, thuB120[P*Cbr-his-72*::cherry] *Cbr*X or thuN14[P*Cbr-his-72*::cherry] *Cni*X. Progenies expressing these reporters were considered as XX females while those without detectable reporter fluorescence were considered as XO males. The *Cbr-his-72* promoter transgene does not express detectable Cherry reporter protein until 100-cell stage. As a result, sexes of embryos younger than the 100-cell stage were undiscernible. Parental strain names of crosses in each figure are listed in Table S4.

Nomenclature of hybrids

The genotypes of the hybrid F1 progeny were indicated by their parents-of-origin in parentheses after F1, which were separated by a comma. The letters "*Cbr*" and "*Cni*" were used to denote C. briggsae and C. nigoni, respectively. For example, F1(♂*Cbr*, ♀*Cni*) represents F1 hybrids from a cross between C. briggsae male and C. nigoni female animals, while F1(♂*Cni*, ♀*Cbr*) represents F1 hybrids from the reciprocal cross. If a parents were introgression-bearing C. nigoni strains (ZZY10330 or ZZY10337), the "*Cni*" in the F1 name was replaced by the strain name. For example, F1(♂*Cbr*, ♀ZZY10330) represents F1 hybrids from a cross between C. briggsae male and ZZY10330 female animals

Determination of male viability

Embryonic arrest of hybrids

Onto preparation plate, we placed 20 L3-stage females on each plate around 30 plates total, either *C. nigoni* or *Cbr-she-1*(v49). After overnight incubation at 25℃, these preparation plates were scrutinized to confirm the females’ non-pregnant status and absence of males. After dubious plates were discarded, these non-pregnant young adult females were transferred to a single 60 mm mating plate which had round 500 adult males. The plate was place into a 25 ℃ incubation for 10 hours for these parent worms to mate. Post-mating, we washed the laid embryos off these plates and applied a 30-second bleaching treatment to these embryos to sacrifice parents.

The embryos were then incubated in M9 buffer at 25 ℃. Embryos that did not hatch after 15-hr incubation were considered inviable. These embryos were centrifuged at 100 rcf for 30 seconds to remove supernatant, followed by suspended in cold methanol containing 1% PFA and 10 ng/mL Hoechst 33342, then rapidly frozen in liquid nitrogen for a minimum of 2 hours. These prepared samples were either immediately mounted on adhesive slides for microscopy or stored long-term at -20 ℃. Nuclei staining by Hoechst was employed to determine developmental stages of unhatched embryos. Embryos displayed fluorescence of the X-linked reporters inherited from their fathers were considered as females, whereas embryos without detectable fluorescence were considered as males.

Hybrid male embryo viability

We quantified male embryo viability by male embryo hatching rate, calculated as the ratio of hatched male embryos to hatched female embryos. In crosses involving *C. nigoni* as maternal parents, L3-stage females were picked up one day before crossing as described in last section. In crosses involving *C. briggsae* as maternal parents, L4 hermaphrodites were picked up one day before crossing. After parents were mixed, mating plates were incubated at 25 ℃. All parents were moved to new mating plates daily. Laid eggs were examined twice a day so that newly hatched L1 larvae could be collected. In crosses involving hermaphrodites as mothers, L1 larvae expressing the autosomal reporter transgene inherited from father were considered as hybrids. The sexes of hybrid L1 larvae were determined according to fluorescence of the X-linked reporters from fathers or according to genotying by single-worm PCR.

The single-worm PCR procedure commenced with Proteinase K digestion: each worm was incubated with 200 µg/mL of the enzyme in Taq buffer at 65°C for a comprehensive 90-minute digestion period, ensuring thorough worm breakdown and DNA release. The reaction was then inactivated by a 15-minute incubation at 95°C. We used 1 µL of this lysate in subsequent PCR reactions. The PCR products of *Cbr*X and *Cni*X templates were designed to have different length. A list of all primers for genotyping assays can be found in Table S3.

Next Generation Sequencing

Single Embryo RNA sequencing

In our single-embryo RNA sequencing study, males carrying an X chromosome reporter were crossed dioecious, non-fluorescent females. Laid embryos were picked and placed on agarose pads. Developmental stages and sexes were determined using a Zeiss A2 fluorescence compound microscope at 100x oil immersion. Selected embryos were then transferred into water with a mouth pipette, undergoing a triple wash prior to single embryo reverse transcription.

For full-length cDNA synthesis, we utilized the Discover-sc WTA Kit V2from Vazyme Biotech (Nanjing, China) (Item #N711-03). Each washed embryo was immediately flash-frozen in liquid nitrogen upon transfer to the Vazyme lysis buffer, supplemented with 1.25 μg/μL QIAGEN Protease (Item #19157)and Oligo dT Primer. These frozen embryos could be stored at -80°C for prolonged periods or immediately processed for lysis and reverse transcription. The lysis process involved thawing embryos on ice and subjecting them to five liquid nitrogen freeze-thaw cycles, followed by a 20-minute incubation at 50°C and a 5-minute deactivation of Qiagen Protease at 85°C. Post-lysis, reverse transcription was performed, and the full-length cDNA was amplified through 17 PCR cycles to ensure optimal results. Subsequent to cDNA synthesis, RNA quality was verified using the Agilent 2100 Bioanalyzer. Embryos showing significant degradation were discarded.

For sequencing library preparation, 5 ng of the high-quality full-length cDNA was tagmented using the TruePrep DNA Library Prep Kit (TD502-02) from Vazyme. This was followed by a PCR amplification with NEBNext® High-Fidelity 2× PCR Master Mix for 5 cycles with custom Nextera dual index primers and purification using 2× VAHTS DNA Clean Beads.

Single Worm genome sequencing

To conduct single-worm DNA sequencing, individual young adult worms were thoroughly washed three times in water and then placed into a lysis buffer. The buffer composition was 5 µL containing 10 mM Tris-HCl (pH 8.3), 50 mM KCl, 0.45% Triton X-100, and 4 µg/µL Qiagen Protease for enzymatic digestion. Each worm was digested at 50°C for 3 hours, with a subsequent heat inactivation of the Qiagen Protease at 70°C for 15 minutes. For tagmentation, 2 µL of 5× TTBL and 0.4 µL of TTE mix V5, components of the TruePrep DNA Library Prep Kit (Item #TD502-02) from Vazyme, were added to the 5 µL lysate. The final volume was adjusted to 10 µL with nuclease-free water. The lysate was pipetted up and down 20 times to ensure homogeneity, followed by a 10-minute incubation at 55°C for Tn5-mediated tagmentation. The reaction was terminated by adding 2.5 µL of 5× TS (TD502-02), and samples were allowed to rest at room temperature for 5 minutes. PCR enrichment was subsequently performed with custom Nextera dual index primers for 15 cycles using the PCR enzyme provided in the Vazyme TruePrep DNA Library Prep Kit (TD502-02). The resulting DNA was purified using 0.9× VAHTS DNA Clean Beads (Vazyme).

The prepared of both RNA-seq and DNA-seq libraries were dispatched to Novogene (Tianjin, China) to use Illumina NovaSeq 6000 for paired-end (2 × 150 bps) sequencing. We obtained more than 30 million reads of high-quality score (greater than 30 mean quality score) per RNA-seq sample and 0.5 million per DNA-seq sample.

Bioinformatics

Reference genome assembly and gene annotation files

Most reference genome and gene annotation files were downloaded from the WormBase or from the NCBI (Table S5). However, the NCBI has the latest genome sequence file of *C. nigoni* JU1421 strain, but no corresponding annotation file. So based on downloaded genome assembly and gene annotation files of *C. nigoni* JU1422 strain, we used Liftoff (v1.6.3) (*12*) to transfer annotation of the JU1422 strain to the JU1421 strain, creating its annotation GFF3 file. The final GTF files for the *C. briggsae* AF16 strain and the *C. nigoni* JU1421 strain were converted from GFF3 using AGAT (v0.8.0)(*13*). At last, their genome assemblies and annotation were merged as metagenome.

Analysis of Next Generation Sequencing (NGS) data

The NGS data were preprocessed by FASTQC (v0.12.1) and Trimmomatic (v0.39) with default parameters to discard low quality reads and trim adapter sequences, respectively.

For the RNA-seq data, we employed HISAT2 for alignment against the metagenome, which encompassed both *C. briggsae* and *C. nigoni* reference genome sequences. Subsequently, gene expression level was obtained using featureCounts (v2.0.6) (*14*) with "-t exon –p -C" parameters based on the combination of *C. briggsae* and *C. nigoni* GTF files. These read counts were converted to counts per million (CPM) to facilitate comparisons across samples. We first compared CPMs of every protein-coding gene in the metagenome between *C. briggsae* samples and *C. nigoni* samples. For 48,251 genes, their ratios of CPMs in the sister species to those in their own species were less than 0.01 (Data S1). These 48,251 genes were used for following analysis to diminish cross-species mapping contamination. To calculate chromosomal expression level, only gene pairs with one-to-one orthology and homologous chromosomal location between *C. briggsae* and *C. nigoni* were taken into account to facilitate interspecific comparison. As a result, 13,603 ortholog gene pairs (Data S2) were used to calculate chromosomal expression level. The CPM of each gene was re-calculated assuming that the metagenome had only these 13,603 pairs of genes.

The DNA-seq data were processed with Bowtie2(*15*) against the metagenome of *C. briggsae* and *C. nigoni*. Next, we partitioned every chromsomes into 1 kb windows. We then utilized featureCounts (v2.0.6)(*14*) to count the number of reads mapped to these 1 kb windows. These read counts were converted to counts per million (CPM), facilitating comparisons across samples. The heatmap of this data were generated by the ComplexHeatmap (v2.16.0) package(*16*) in R (v4.3.1).

Identifying homologs and updating reference genome

For the 16 genes implicated in the dosage compensation and sex determination pathway, we utilized the Gene Tree feature of WormBase ParaSite (parasite.wormbase.org) (*17*) to find their orthologs and paralogs across *Caenorhabditis*. If a query gene had multiple orthologs in a species, their synteny with the query gene was examined to confirm orthology. The graphics of synteny were generated by genoplotR(*18*). Moreover, we examined genes flanking the query to resolve putative gene complex due to duplication, deletion and or pseudogenization. According to the orthologous and paralogous relationships, we assigned new gene names to seven sequence names in this study, such as a *C. tribulationis* gene (*Ctr-xol-1* for *CSP40.g8311*) and four *C. nigoni* genes (*Cni-xol-1* for *Cnig_chr_X.g24950*, *Cni-xol-1.1* for *Cnig_chr_X.g24897*, *Cni-bnsp-1* for *Cnig_chr_II.g7047*, and *Cni-bnsp-2* for *Cnig_chr_II.g7048*). The WormBase ParaSite suggested that neighbor genes *Cni-bnsp-1* and *Cni-bnsp-2* had a 2-to-1 ortholog in *C. briggsae*, *CBG19535*. In the *C. briggsae* genome, *CBG19535* and pseudogene *CBG19537* are homologous neighbors in a such a way that pseudogene *CBG19537* has a syntenic location with *Cni-bnsp-1*. Moreover, *CBG19537* is the reciprocal best blastn hit of *Cni-bnsp-1*, suggesting their orthology (*19*). Therefore, *CBG19537* and *CBG19535* were assigned gene names *Cbr-bnsp-1* and *Cbr-bnsp-2*, respectively.

Noticeably, the gene structure of *Cbr-bnsp-2* in the WormBase has a 49-bp intron that does not exist in *Cni-bnsp-2*. But according to our RNA-seq reads, this intron should be part of its flanking exons. Moreover, our RNA-seq reads also revealed an 1bp insertion 'T' in exon 6 due to assembly error. We updated the annotation and sequence of *Cbr-bnsp-2* correspondingly(*8*).

In the downloaded genome assembly of *C. tribulationis*, scaffold CSP40.scaffold00353, harboring *Ctr-xol-1*, and CSP40.scaffold01719, exhibit overlapping sequences. Furthermore, these scaffolds demonstrate collinearity with the continuous *Cni*X region, which also harbors *Cni-xol-1*. Consequently, CSP40.scaffold00353 and CSP40.scaffold01719 are highly likely to represent consecutive segments of the *Ctr*X region. Based on BLASTn end alignments, scaffolds CSP40.scaffold00353 and CSP40.scaffold01719 were merged and subsequently polished with Pilon (*20*), resulting in the formation of a new scaffold named 'merge-CSP40.scaffold00353-CSP40.scaffold01719_pilon'(*8*). The annotations from the two original scaffolds were transferred to the new scaffold using Liftoff (v1.6.3) (*12*).

Phylogenetic analysis

The CDS of *Cbr-xol-1*, and *Ctr-xol-1* were downloaded from the WormBase. *Cni-xol-1*, *Cni-xol-1.1* were extracted from the downloaded genome assembly of the JU1421 strain (Table S5). Codon-based nucleotide alignment of these CDS was created using the ClustalW (codon) feature in MEGA11(*21*). Phylogenetic analyses of these *xol-1* genes were conducted using the maximum likelihood method implemented in MEGA11(*21*). Our nucleotide-based phylogeny used the General Time Reversible model. We inferred the bootstrap consensus tree from 1000 replicates. Initial tree(s) for the heuristic search were obtained automatically by applying Neighbor-Joining and BioNJ algorithms. A discrete Gamma distribution was used to model evolutionary rate differences among sites (five categories). Only the 3rd codon position was selected for analysis. We chose to remove all sites containing alignment gaps and missing information (Complete-deletion option).

smiFISH and transcript counting

The smiFISH was performed as described (*22*) with modification. The probes used in this study are designed at Stellaris Probe Designer (<https://www.biosearchtech.com/stellaris-designer>). A set of Cherry probes were used to detect whether scored embryos carried X-linked Cherry reporter inherited from father so that the sex of embryos could be determined. A set of rRNA probes were used as hybridization quality control. For each probe set, the primary probes were synthesized by Azenta Life Science (Tianjin, China). The secondary fluorescently-labeled FLAP probes were synthesized either by Sangon Biotech (Shanghai, China) or by Azenta Life Science (Tianjin, China). Information on these probes can be found at Data S3. The probes were solubilized in TE buffer at concentration of 100 mM. The primary probes were combined at equimolar ratio. Than the primary probes set were mixed with secondary fluorescently-labeled FLAP probe at molar ratio of 4:5 in 1x NEB3/3.1 buffer so that the concentration of each primary probe and the secondary probe were 20 uM and 25 uM, respectively. These probes were annealed at 95 °C for 3 min, followd by decreasing the temperature at 0.1 °C/min to 45°C and incubating for 2 min at each 10 °C interval.

In each cross, a mix of 600 adult males and 600 young adult females was placed on a 60 mm NGM plate and then incubated at 25 ℃ for 17 hours. Subsequently, the adults were transferred to a new plate to continue laying embryos for additional 12 hours to increase the number of embryo samples. Laid embryos in these plates were washed off, subjected to a 30 to 45-second bleach treatment, and then rinsed three times to remove bleaching buffer. After supernatant was removed, 1 mL of -20 ℃ pre-chilled anhydrous methanol was added to embryos, followed by flash-freezing in liquid nitrogen for at least 12 hours. Subsequently, these embryos were centrifuged to remove methanol, and then fixed in 1% PFA at room temperature for 10 minutes. After PFA was removed, the samples were washed with 1x PBS 3 times and then treated overnight in 70% ethanol at 4°C. After removing the ethanol, the embryos were processed with 2x SSC solution containing 0.5% Triton X-100 for 10 minutes, followed by pre-treatment for one minute with 30% formamide Wash Buffer A (200 μL Stellaris Wash Buffer A <Biosearch Technologies, SMF-WA1-60>, 300 μL deionized formamide <Millipore, S4117>, and 500 μL DEPC treated RNAse free water <Invitrogen, AM9922>).

The 30 % formamide pretreated embryos were suspended in hybridization buffer (99 μL Stellaris Hybridization buffer <Biosearch Technologies, SMF-HB1-10), 11 μL deionized formamide). Then 6 μL of pre-annealed probes (2 μL of target gene, 2 μL of Cherry, and 2 μL of rRNA) were added to the sample and the mixture was placed in a 37 ℃ incubator for 8 to 48 hours for hybridization. The samples were then rinsed with 10% formamide Wash Buffer A (200 μL Stellaris Wash Buffer A, 100 μL deionized formamide and 700 μL DEPC treated RNAse-free water) for 45 minutes. After removing Wash buffer A, the samples were stained with Hoechst in 10% formamide Wash Buffer A for 45 min. Next, these embryos were washed three times with Stellaris Wash Buffer B (Biosearch Technologies, SMF-WB1-20), 5 min each, then resuspended in 15-20 µL of mounting medium (50% glycerol <Sigma, G5516-100 mL>, 2% N-progl gallate <Sigma, 02370-100 g> and 50 mM Tris-HCL pH 8.0) and incubated at 4°C for 30 minutes before confocal microscopy imaging. The genotypes of all parental strains used for smiFISH are detailed in the Table S4.

Images were captured using a Zeiss 780 inverted confocal microscope, outfitted with a Plan-Apochromat 63x/1.40 Oil DIC M27 objective. The acquisition parameters were as follows: a 512x512 frame size, Zoom 2.5, and a z-interval of 0.2 µm. For the Quasar 570 dye, a 561 nm laser was set to 90% power, with the pinhole adjusted to 1 Airy Unit (AU), and the detection range was 561-636 nm. The gain was set to 800. For the Hoechst dye, excitation was achieved using a 405 nm laser with the pinhole set to 1 Airy Unit (AU). The laser power was configured at 2%. The gain was adjusted as needed based on the image feedback to prevent overexposure and maintain the quality of the fluorescence signal. Alexa fluor 488 and Cy5 were acquisition with a dual-color channels, the 488 nm and 633 nm lasers with 80% laser power were employed. The pinhole was set to 1 Airy Unit (AU), the gain was set to 800 for both channels. Additionally, all acquisition process incorporated a 4 times line-wise summation.

Tiff stack images from confocal imaging were processed using custom ImageJ macro, a robust platform for image analysis. For spot detection, we utilized the Big-FISH package(*23*), specifically designed for the precise identification of single-molecule FISH signals. To determine the subcellular localization (nucleus or the cytoplasm) of each detected punctum, we converted the spot data of each embryo into VANO-compatible formats using custom ImageJ macro. The precise location of each spot within individual embryos was then manually curated using VANO(*24*).

In addition, to accurately assess the embryonic stage, we employed the 'Cell_Segmentation_GVF' plugin within the Vaa3D software(*24*). This enable us to estimate the cell stage based on the segmentation results.

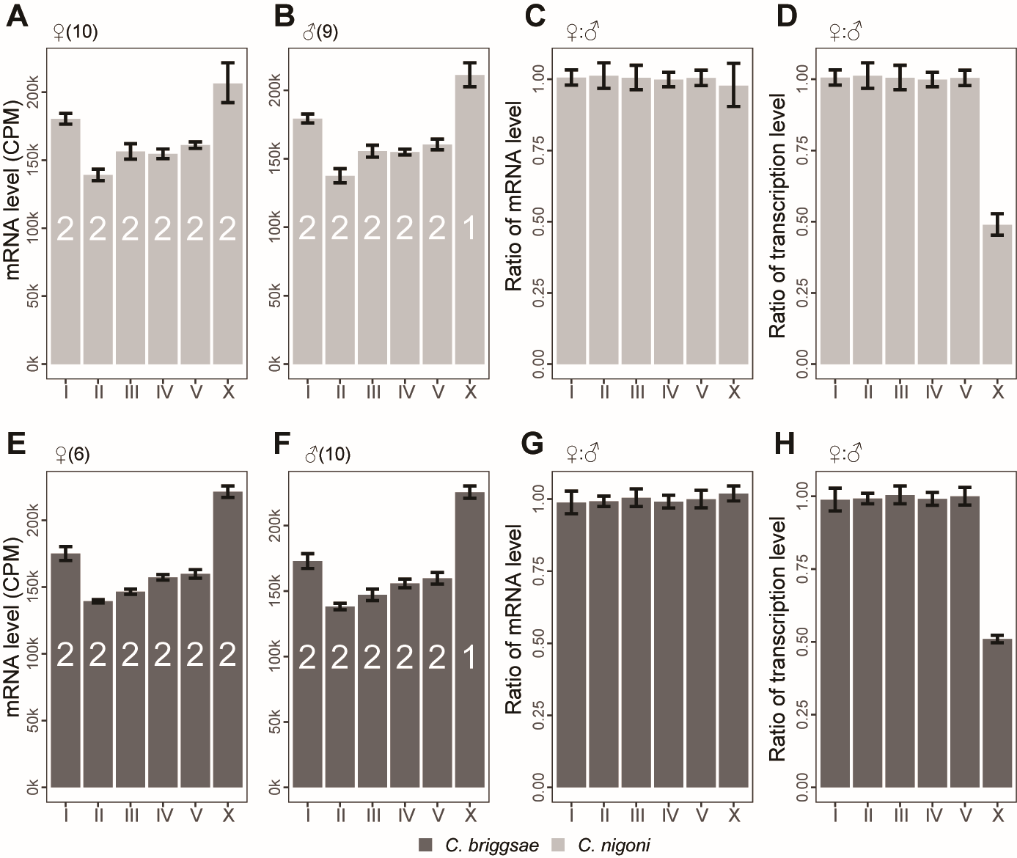

fig. S1. Balanced ChrX expression between sexes in both parental species.

(**A** and **B**) Expression levels of chromosomes in female (A) and male (B) embryos of *C. nigoni*. The copy number of every chromosome is denoted by a white numeral. The number of profiled embryos is indicated in parentheses. (**C** and **D**) Comparison of chromosomal expression levels (C) and transcription levels (D) between two sexes of *C. nigoni*. The transcription level is defined as the mRNA level divided by the copy number of chromosome. (**E** and **F**) Expression levels of chromosomes in female (E) and male (F) embryos of *C. briggsae*. Their mothers were *she-1*(*vp49*)/ *she-1*(*vp49*) so that no embryo was derived from selfing. Labels are as in (A) and (B). (**G** and **H**) Comparison of chromosomal expression levels (G) and transcription levels (H) between two sexes of *C. briggsae*. The color codes for species origin of chromosomes are as indicated. error bar, 95% confidence interval.

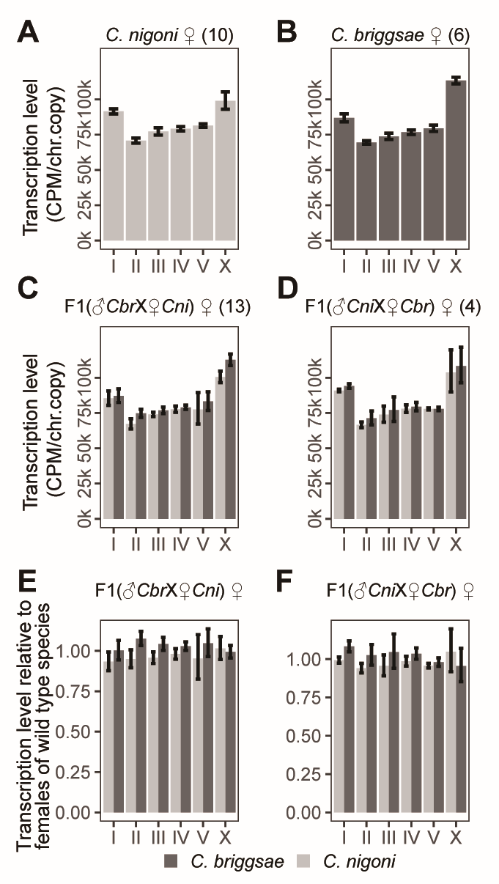

fig. S2. Dosage compensation in hybrid female embryos.

(**A** to **D**) Transcription levels of chromosomes in *C. nigoni* females (A), *C. briggsae* females (B), hybrid females from *C. nigoni* mothers (C), and hybrid females from the reciprocal cross (D). The number of profiled embryos is indicated in parentheses. (**E** and **F**) Comparison of chromosomal transcription levels in females between hybrids and their parental species. The color codes for species origin of chromosomes are as indicated. error bar, 95% confidence interval.

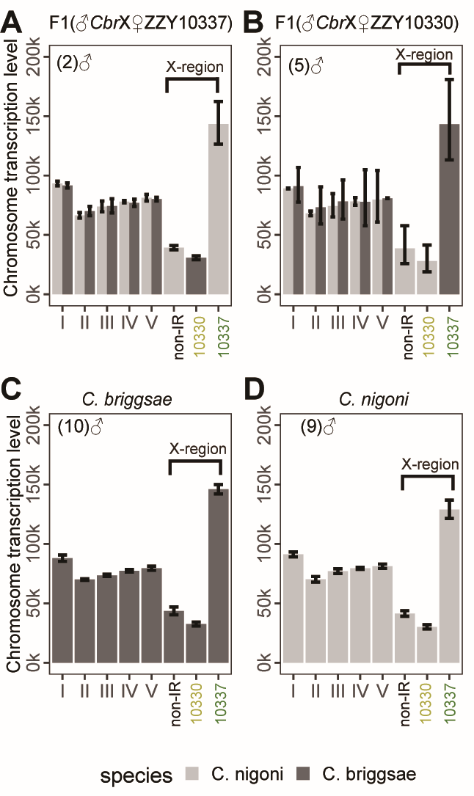

fig. S3. Chromosome transcription levels in male embryos with computational division of ChrX into three regions defined in Fig. 2A

(**A** and **B**) Hybrids male embryos from ZZY10337 mother (A) and from ZZY10330 mother (B). (**C** and **D**) Male embryos of *C. briggsae* (C) and *C. nigoni* (D). There is no data on cross involving either ZZY10337 or ZZY10330 as paternal parent because these two HIL strains are male sterile(*25*). The color codes for species origin of chromosomes are as indicated. error bar, 95% confidence interval.

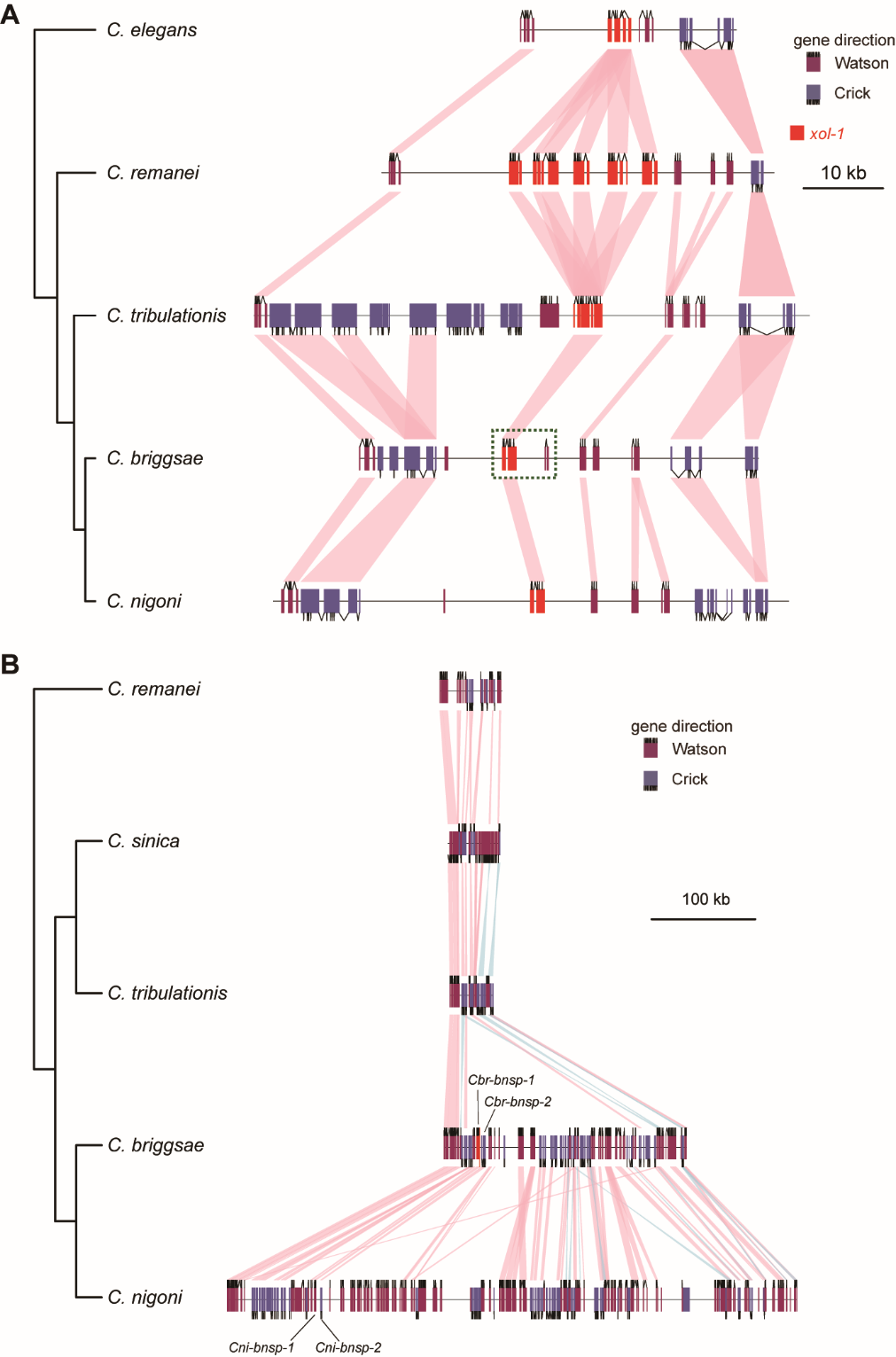

fig. S4. Evolution of two genes regulating dosage compensation.

(**A**) Collinear blocks where *xol-1* orthologs reside across five *Caenorhabditis* species. The *Cbr*X segment within green dashed frame are compared to a *Cni*X segment harboring *Cni-xol-1.1* in Fig. 3E. (**B**) ChrII segments where *bnsp-1 and bnsp-2* resides and their orthologous blocks in other three *Caenorhabditis* species. Pink pastel shapes connect orthologs with same orientation, while blue ones connect inverted orthologs.

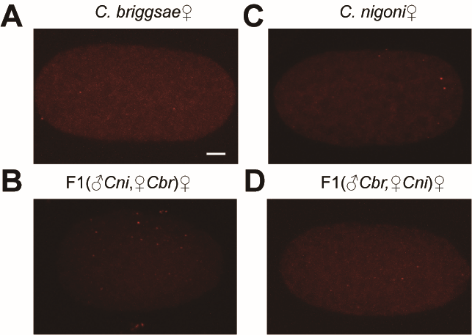

fig. S5. The mRNA molecules of *xol-1* genes detected in female embryos by smiFISH.

**(A** and **B)** *Cbr-xol-1* mRNA molecules in female embryos of *wild-type* (A) and hybrids sired by *wild-type* *C. nigoni* (B). Probes used in these embryos are designed for *Cbr-xol-1.* **(C** to **D)** *Cni-xol-1* and *Cni-xol-1.1* mRNA molecules in female embryos of *wild-type* *C. nigoni* (C) and hybrids sired by wild type *C. briggsae* (D). Probes used in these embryos are designed for both *C. nigoni xol-1* genes so that a punctum can be either *Cni-xol-1* mRNA or *Cni-xol-1.1* mRNA. Every embryo is oriented so that its anterior region is to the left. The number of scored embryos is indicated in parentheses. Embryos were collected from the 500-cell to the bean stages. Scale Bar 5 um.

Table S1. Strains in this study

| **Strain** | **Species** | **Genotype** |
| --- | --- | --- |
| AF16 | *C. briggsae* | *Wild-type* |
| Cbr-she-1 | *C. briggsae* | *Cbr-she-1(v49)* |
| JU1421 | *C. nigoni* | *Wild-type* |
| XIL1391 | *C. briggsae* | *thuExB391[PCbr-xol-1(8.9kb)::Cbr-xol-1::Cbr-xol-1 3'UTR;pZZ184(PCbr-myo-2::mcherry::Prps-27::NeoR)];Cbr-she-1(v49), IV* |
| XIL31003 | *C. briggsae* | *Cbr-xol-1(thuB124),line1, X;thuB134(PCni-xol-1.1::Cni-xol-1.1::Cni-xol-1.1 3'UTR), II* |
| XIL31006 | *C. briggsae* | *thuB129(PCni-xol-1(9.7kb)::Cni-xol-1::Cni-xol-1 3'UTR), II;Cbr-xol-1(thuB124),line1, X;thuB134(PCni-xol-1.1::Cni-xol-1.1::Cni-xol-1.1 3'UTR), II* |
| XIL31008 | *C. briggsae* | *thuB119(PCbr-his-72::wCherry::H2B::Cbr-his-72 3'UTR X);Cbr-she-1(v49) IV;thuB135(replace CBG19537 gene region by Cnig_chr_II.g7047(Cni-bnsp-1)), II* |
| XIL3957 | *C. briggsae* | *thuB119(PCbr-his-72::wCherry::H2B::Cbr-his-72 3'UTR X);Cbr-she-1(v49) IV* |
| XIL3960 | *C. nigoni* | *thuN26(PCbr-his-72::myri::cherry::Cbr-his-72-3'UTR), II;thuN14(PCbr-his-72::wCherry::H2B::Cbr-his-72-3'UTR), X;* |
| XIL3963 | *C. briggsae* | *thuB119(PCbr-his-72::wCherry_H2B::CB-his-72 3'UTR), X;Cbr-she-1(v49), IV;thuB120(PCbr-his-72::myri_cherry::Cbr-his-72 UTR), II* |
| XIL3985 | *C. briggsae* | *thuB129(PCni-xol-1(9.7kb)::Cni-xol-1::Cni-xol-1 3'UTR), II;Cbr-xol-1(thuB124),line1, X；* |
| XIL6118 | *C. briggsae* | *thuB118(PCbr-his-72::neonGreen::H2B::Cbr-his-72 3'UTR), X* |
| XIL6124 | *C. briggsae* | *Cbr-xol-1(thuB124),line1, X* |
| XIL6129 | *C. briggsae* | *thuB129(PCni-xol-1(9.7kb)::Cni-xol-1::Cni-xol-1 3'UTR), II* |
| XIL6135 | *C. briggsae* | *thuB135(replace CBG19537 gene region by Cnig_chr_II.g7047(Cni-bnsp-1)), II* |
| XIL714 | *C. nigoni* | *thuN14(PCbr-his-72::wCherry::H2B::Cbr-his-72 3'UTR), X* |
| XIL723 | *C. nigoni* | *Cni-xol-1.1(thuN23);thuN14(PCbr-his-72::wCherry::H2B::Cbr-his-72-3'UTR), X* |
| XIL3999 | *C. nigoni* | *Cnig_chr_II.g7047(Cni-bnsp-1)(thuN29),II;thuN14(PCbr-his-72::mCherry::H2B::Cbr-his-72 3' UTR), X* |
| XIL729 | *C. nigoni* | *Cnig_chr_II.g7047(Cni-bnsp-1)(thuN29),II* |
| XIL734 | *C. nigoni* | *Cni-xol-1(thuN24), X/+;thuN30(Cni-xol-1[PCbr-his-72::neonGreen::H2B::Cbr-his-72 3'UTR::Prps-27::NeoR/KanR::unc-54 3'UTR]). X/+* |
| ZZY10330 | *C. nigoni* | *zzyIR10330[(Cbr-Pmyo-2::gfp),X:16.36 Mbp-21.54 Mbp,AF16]* |
| ZZY10337 | *C. nigoni* | *zzyIR10337[Cbr-unc-119(+); myo-2::GFP,X:3.34 Mbp-14.62 Mbp,AF16]* |
| XIL6104 | *C. briggsae* | *Cbr-she-1(v49), IV;Cbr-him-8(v188) I; stIs20120 [PCbr-myo-2::GFP + Cbr-unc-119(+)], X* |
| XIL752 | *C. nigoni* | *Cni-xol-1.1(thuN52), X/+;thuN53[(Cni-xol-1/+,Cni-xol-1(PCbr-his-72::neonGreen::H2B::Cbr-his-72 3'UTR::Prps-27::NeoR/KanR::unc-54 3'UTR)], X* |
| XIL31009 | *C. briggsae* | *thuB134(PCni-xol-1.1::Cni-xol-1.1::Cni-xol-1.1 3'UTR), II;cbr-she-1(v49), IV* |

Table S2. CRISPR-Cas9 Targets in this study

| **gRNA** | **Purpose** |
| --- | --- |
| cgtggggaaagtctgcgccc | *thuB118* |
| cgtggggaaagtctgcgccc | *thuB119* |
| gctctataaaggcaccgcgg | *thuB120* |
| ccatatcgcaaacgatggct,tctcgcgcccctccagggta,gacgttggagataacgcgcc,agacggtgactcaactctgg | *thuB124* |
| gctctataaaggcaccgcgg | *thuB129* |
| CGACACCCTAAACCCTGCCA,GCATAAGGTCGATTATCCAT | *thuB134* |
| ATTCCTACGAATGAAGAAAG,TAGAGCTTCTTGGAAGACGA | *thuB135* |
| CGTGGGGGAAGTCTGCGCCC | *thuN14* |
| GTACTGTAAAGCTACCACCA,AAAGTCGTTAGAGCATCCAG,CGCAGAGACATTCTACGCAA | *thuN23* |
| GTACTGTAAAGCTACCACCA,AAAGTCGTTAGAGCATCCAG,CGCAGAGACATTCTACGCAA | *thuN24* |
| gctctataaaggcaccgcgg | *thuN26* |
| CAGGACACCAATACTCCGGC,GATGGGATGATTCTGCAAGA | *thuN29* |
| CGCAGAGACATTCTACGCAA | *thuN30* |
| GTACTGTAAAGCTACCACCA,CGCAGAGACATTCTACGCAA,CTCTTCCAATTGAAGAGTAA | *thuN52* |
| GTACTGTAAAGCTACCACCA,CGCAGAGACATTCTACGCAA,CTCTTCCAATTGAAGAGTAA | *thuN53* |
| attcgcgtcagatgatgtac | co-CRISPR |

Table S3. Primers in this study

| forward | reverse | PCR product size(bp) | Purpose | Source |
| --- | --- | --- | --- | --- |
| attttcgtttgatttggtcaaaaacca | cagctgtccctctcattccaaa | WT:1475 KI:5560 | *thuB118* genotyping | This study |
| attttcgtttgatttggtcaaaaacca | cagctgtccctctcattccaaa | WT:1475 KI:5509 | *thuB119* genotyping | This study |
| tgtggagaatgttgggttca | acaccttcgaccctcctttt | WT:1574 KI:5263 | *thuB120* genotyping | This study |
| TTGTCCTCGAGGACTTCACTATTTC | TGGCTGGACTATTCTTTTGTATTCACC | WT:3027 KO:2563 | *thuB124* genotyping | This study |
| tgtggagaatgttgggttca | TCAGTGGTATGGTTTTTCTTAGCG | 1458 | Left External Primer and Internal Primer for *thuB129* | This study |
| CACGCAACCGTCAGTATTAG | TGCTATGGAGAACAAAGTCAAGT | 1909 | Right External Primer and Internal Primer for *thuB129* | This study |
| TGTGTACGTAGACAGGCCTGG | AAGCGCTCTGTAGTCCATGAGT | 1697 | Left External Primer and Internal Primer for *thuB134* | This study |
| CAATTCCCCTTCGGTCAAGCTT | TCAGCGCCAAATTCTCGAAGC | 1696 | Right External Primer and Internal Primer for *thuB134* | This study |
| AGCCAACTTGAGGTGATTTGAT | TACACACTGACTAGAGTAGACGG | 1496 | *thuB135* genotyping | This study |
| GTTCCTCTCAGAGTCAACTCTCT | TGTGCTGCACAATGTATCTTCT | 1451 | *thuB135* genotyping | This study |
| cctccagggtaggggtaga | cgcaaacgcaaaatcgtaattg | WT:1260 KO:614 | *thuN23* genotyping | This study |
| CTGCATGTGTCAAAACGGGA | GTTCGAATCTTGCTTGTGGATTTC | WT: 2763 KO:2014 | *thuN24* genotyping | This study |
| ATGGCTCCAGAAATGGATTGG | ATTCATCTGGATCAACTAACATCCG | WT:1453 KI:5142 | *thuN26* genotyping | This study |
| TCTGCACTCTCTTCGTTCTCAA | CTGCCAACTATCCTTAGGATCTCA | WT:3261 KO:1380 | *thuN29* genotyping | This study |
| CTGCATGTGTCAAAACGGGA | aaatagcctgaaaactgaaaattttgaaaattc | 2594 | Left External Primer and Internal Primer for *thuN30* | This study |
| TCAACATCCCTACATGCTCTTTCT | AACCTGGATATTCTAGAAAAATAGGGATTAAATT | 1729 | Right External Primer and Internal Primer for *thuN30* | This study |
| CTGCATGTGTCAAAACGGGA | AACCTGGATATTCTAGAAAAATAGGGATTAAATT | WT:3501 KI:9891 | External Primer for *thuN30* | This study |
| TCACTGGACTCATCGACTGCTGGAATC | CAAATGGAGGAGGGAGGAGAAAATGCG | C. nigoni:552;  C. briggsae:593 | Nest Outer Primer for *vab-3* loci, Distinguish chrX origin | This study |
| tgcactcgggcatactgtaa | tgtacaacgggctcagtcag | *C. nigoni*:269;  *C. briggsae*:334 | Nest Inner Primer for *vab-3* loci, Distinguish chrX origin | (*26*) |
| ctcactgttcatagaaggccgtatccg | gaccgagctcctccacctctactattag | C. nigoni:350;  C. briggsae:573 | Nest Outer Primer for *dpy-21* loci, Distinguish chrV origin | This study |
| ATCCGATGGTGGTGGTAGTG | CCGCAGCGCAGTAAGTTTT | *C. nigoni*:283;  *C. briggsae*:506 | Nest Inner Primer for *dpy-21* loci, Distinguish chrV origin | This study |

Table S4. Parental strain names of crosses for each figure

| **father strain** | **mother strain** | **figure** |
| --- | --- | --- |
| XIL3957 | JU1421 | Fig. 5B |
| XIL714 | JU1421 | Fig. 5A |
| XIL3957 | Cbr-she-1 | Fig. 5D |
| XIL31008 | XIL6135 | Fig. 5E |
| XIL714 | Cbr-she-1 | Fig. 5F |
| XIL3999 | XIL729 | Fig. 5G |
| XIL714 | JU1421 | Fig. 3B |
| XIL714 | XIL752 | Fig. 3B |
| XIL714 | XIL734 | Fig. 3B |
| JU421 | XIL723 | Fig. 3B |
| XIL1391 | XIL734 | Fig. 3B |
| XIL3963 | XIL6124 | Fig. 3C |
| XIL6129 | XIL6124 | Fig. 3C |
| XIL3963 | XIL31003 | Fig. 3C |
| XIL31008 | XIL6135 | Fig. 6E |
| XIL3957 | AF16 | Fig. 6E, Fig. 3C |
| XIL714 | Cbr-she-1 | Fig. 5I |
| XIL714 | Cbr-she-1 | Fig. 6E, Fig. 5I,  Fig. 4A |
| XIL714 | XIL31012 | Fig4. A |
| XIL3999 | Cbr-she-1 | Fig. 6E |
| XIL714 | Cbr-she-1 | Fig. 5I |
| XIL714 | Cbr-she-1 | Fig. 5I |
| XIL3960 | XIL3985 | Fig. 4A |
| XIL3960 | XIL31006 | Fig. 4A |
| XIL3960 | XIL31003 | Fig. 4A |
| XIL3957 | JU1421 | Fig. 4A |

Table S5. Downloaded Genome and annotation files in this study

| **species** | **Strain** | **file type** | **file name** | **source website** |
| --- | --- | --- | --- | --- |
| *C. briggsae* | AF16 | genome assembly | c_briggsae.PRJNA10731.WS290.genomic.fa.gz | [www.wormbase.org](http://www.wormbase.org) |
|  |  | annotation | c_briggsae.PRJNA10731.WS290.annotations.gff3.gz | [www.wormbase.org](http://www.wormbase.org) |
|  |  | ortholog | c_briggsae.PRJNA10731.WS290.orthologs.txt.gz | [www.wormbase.org](http://www.wormbase.org) |
| *C. elegans* | N2 | annotation | c_elegans.PRJNA13758.WS290.annotations.gff3.gz | [www.wormbase.org](http://www.wormbase.org) |
|  |  | ortholog | c_elegans.PRJNA13758.WS290.orthologs.txt.gz | [www.wormbase.org](http://www.wormbase.org) |
| *C. nigoni* | JU1422 | genome assembly | c_nigoni.PRJNA384657.WS290.genomic.fa.gz | [www.wormbase.org](http://www.wormbase.org) |
|  |  | annotation | c_nigoni.PRJNA384657.WS290.annotations.gff3.gz | [www.wormbase.org](http://www.wormbase.org) |
|  | JU1421 | genome assembly | GCA_027920645.1_ASM2792064v1 | <https://www.ncbi.nlm.nih.gov/assembly/GCA_027920645>.1 |
| *C. remanei* | PX506 | annotation | c_remanei.PRJNA577507.WS290.annotations.gff3.gz | [www.wormbase.org](http://www.wormbase.org) |
| *C. sinica* | JU800 | annotation | c_sinica.PRJNA194557.WS290.annotations.gff3.gz | [www.wormbase.org](http://www.wormbase.org) |
| *C. tribulationis* | JU2818 | annotation | c_tribulationis.PRJEB12608.WS290.annotations.gff3.gz | [www.wormbase.org](http://www.wormbase.org) |
|  |  | genome assembly | c_tribulationis.PRJEB12608.WS290.genomic.fa.gz | [www.wormbase.org](http://www.wormbase.org) |
