## Supplementary Data S1 to S3 for "Regulatory divergences in dosage compensation cause hybrid male inviability in *Caenorhabditis*": Data_captions.docx

**Data S1. (separate file)**

*C. briggsae* and *C. nigoni* genes whose RNA-seq reads have less than 1% interspecific mapping contamination.

**Data S2. (separate file)**

*C. briggsae* and *C. nigoni* gene pairs with 1:1 ortholog relationship and no inter-chromosomal translocation

**Data S3. (separate file)**

smiFISH probes for *Cbr-xol-1*, *Cni-xol-1*/*1.1*, cherry and rRNA
